# A multisensory, bidirectional, valence encoder guides behavioral decisions

**DOI:** 10.1101/2025.09.26.678749

**Authors:** Claire Eschbach, Katrin Vogt, Bruno Afonso, Nick Polizos, Stephanie Dancausse, Anastasiia Evans, Anna Verbe, Kun Wang, Matthew Berck, Aravi Samuel, Mason Klein, Albert Cardona, Marta Zlatic

## Abstract

A key function of the brain is to categorize sensory cues as repulsive or attractive and respond accordingly. While we have some understanding of how sensory information is processed in the sensory periphery, the classification of cues according to valence in central brain circuits is less well understood. Here, we addressed this question in the *Drosophila* larva, where we could leverage the synaptic resolution connectome to determine where innate and learnt information from distinct aversive and appetitive sensory modalities converges; and combine this with imaging and manipulation of neural activity to determine how valence information is encoded and used to guide navigation. We found that information from multiple innately aversive and attractive sensory modalities converges onto a common output neuron of the learning circuit, specifically the Mushroom Body output neuron, MBON-m1. We discovered that this neuron is required for navigating both attractive odor and aversive temperature cues and is activated by attractive cues, such as food odour and sugar, and inhibited by different aversive cues, such as cooling, salt, or non-food odors. Together, our study reveals a neuron that bi-directionally encodes valence and controls actions.

## Introduction

Natural environments are complex, multisensory worlds from which animals must extract useful information. A key function of the brain is to categorize sensory cues according to their valence (as repulsive or attractive) and respond accordingly. How valence is computed independent of the type of sensory stimulus, encoded, and used to drive approach or avoidance is unclear.

In vertebrates and evolutionarily complex invertebrates (e.g. insects), higher-order associative learning circuits compute the learnt valence of stimuli. At the same time, many stimuli have an innate valence, acquired through evolution, which has to be integrated with the learnt one. While recent studies have started to reveal how the innate and learnt valence of a single stimulus is integrated, how the valence of different sensory modalities is integrated and used to drive behaviour is poorly understood. For example, parallel, modality-specific pathways could converge late in the sensory processing hierarchy, at the descending or pre-descending neurons. Alternatively, a coherent valence signal could be computed in higher-order brain areas based on innate and learnt information from multiple modalities. It is also unclear how positive and negative valence signals are encoded and how conflict between them is resolved. In principle, activation of some neurons could encode positive, and of others, negative valence. Alternatively, the same neurons could bi-directionally encode positive and negative valence.

To address these questions, it is essential to 1) know where and how information from distinct sensory modalities that encode either the same or opposite valence converges in the nervous system and 2) functionally test key convergence nodes. This is challenging in larger, vertebrate nervous systems because we lack information about the synaptic resolution architecture and, hence, the precise patterns of convergence between sensory modalities. Furthermore, it is difficult to selectively record from, and manipulate, uniquely identified convergence nodes. Here, we used the tractable *Drosophila* larva as a model system to study where and how valence signals from distinct sensory modalities are encoded, integrated, and used to guide navigation behaviour.

*Drosophila* larvae navigate a rich sensory world in which most of the perceived sensory stimuli have an innate valence. Thus, larvae innately approach most odours, such as the fruit compound ethyl acetate [1], but avoid a few of them, such as geranyl acetate, an antiherbivore plant compound [2]. They avoid temperatures below 22°C to maintain their body at a homeostasis temperature around 24°C [3]. The gustatory system allows them to find palatable food: they approach sweet substrates [4] and avoid highly salty ones [5,6].

When navigating a sensory gradient, approach or avoidance behaviors are a direct result of the frequency of direction changes. Specifically, an increase in the intensity of an attractive cues (e.g. attractive odour) decreases the probability of turning [7–10], allowing the larva to crawl up the gradient, whereas a decrease in the intensity of an attractive cue increases the probability of turning. Conversely, an increase in the intensity of an aversive cue (such as aversive odour, or temperature), increases the probability of turning, allowing the larva to crawl down the gradient [11–14]. Thus, turn rate, and to a lesser extent turn direction and turn size, are behavioral components used by larvae to navigate gradients of distinct cues [10,11,13–16]. Likewise, these behavioral components are impacted when innate attraction for an odor is shifted towards aversion following the formation of aversive olfactory memory [17].

We already have a good understanding of basic olfactory processing circuits that are used to direct behavioral decisions. In the larval olfactory system, which is similar to the mammalian olfactory system, olfactory receptor neurons (ORNs) project onto specific antennal lobe glomeruli. Different projection neurons (PNs) then convey the olfactory information to two higher-order brain structures, the lateral horn (LH) and the mushroom body (MB), thought to transform odor-quality signals into valence signals [18–20]. The larval MB intrinsic neurons, the Kenyon Cells (KCs), receive mostly olfactory input, but a subset receives visual, thermosensory, and/or other sensory input [21]. All KCs converge onto 24 left-right pairs of MB output neurons (MBONs) that encode learned valences of stimuli (positive or negative) [17,21]. The synaptic connections between KCs and MBONs are subject to experience-dependent dopaminergic-mediated plasticity [18,22]. The LH relays odor signals without any known associative learning-dependent plasticity [23]. We have recently shown that olfactory LH and MBON pathways converge onto specific circuit nodes, such as the MBON-m1. MBON-m1 receives excitatory inputs from olfactory LH pathways that encode positive valence, as well as from MBONs that encode positive valence, and inhibitory input from MBONs that encode negative valence [17]. Consistent with this, we found that the attractive odor activates MBON-m1, and its odor response is decreased after aversive training to the odor. We suggested that this enables the final navigational decision to result from an integration of innate and learnt valence of the cue, even if they are contradictory.

Is innate attraction or aversion towards cues from different sensory modalities also processed via similar convergence circuits? To answer this question, we asked where pathways for innate aversion to cool temperatures converge with pathways for innate attraction to an odor, and how this integrated valence is used to guide navigation.

When navigating temperature gradients with a cold region, the increase in the intensity of an aversive cue (*e*.*g*. decrease in temperature below 22°C) increases the probability of turning, whereas a decrease in the intensity of an aversive cue (*e*.*g*. increase in temperature above 22°C) decreases turn probability [3,11]. Even though larvae use the same behavioral strategy to navigate temperature and olfactory gradients, less is known about the cold processing pathway in larvae. According to EM reconstruction, three sensory Cooling Cells (CCs) closely follow the antennal nerve tract formed by the ORNs from the dorsal organ ganglion (DOG) to the antennal lobe [11,12] and project in the region posterior-dorsal to it. From there, cold thermosensory projection neurons (ctPNs) provide input to the MB calyx onto two thermo-specific KCs [21] and other previously uncharacterized neurons in the LH. The detailed anatomy, connectivity and functions of these thermosensory projection neurons have not been described. The CCs, which receive reciprocal cross-inhibition from Warming Cells, are necessary for cold avoidance [11,24] and confer to the larvae the ability to maintain themselves in their preferred temperature around 24°C. Optogenetic activation of the CCs induces aversive turning in larvae [12].

In this study, we used the connectome to analyse the cold thermosensory pathways and to understand if and where attractive and aversive sensory cues converge in the brain to regulate the same behavioral outputs. We describe and functionally test ctPNs involved in avoiding cooling temperature. Most of their direct postsynaptic partners are KCs or LH neurons. Further downstream, we identified the MB output neuron m1 (MBON-m1) as one of the earliest points of convergence between thermosensory and olfactory pathways. Previously, we have shown that this neuron receives convergent input from olfactory LH and olfactory KC pathways and is activated by innately attractive odours. Here, we find that MBON-m1 also receives convergent input from thermosensory LH pathways, thermosensory KC pathways and multi-sensory PNs, and is inhibited by innately aversive thermosensory inputs. We show that this neuron is required for both odour approach and cold avoidance. We further tested the response of MBON-m1 to a range of different sensory stimuli and provide evidence that MBON-m1 is a multimodal neuron that responds to sensory stimuli according to their valence: it is inhibited by innately aversive stimuli, including cold, aversive odours and taste (salt), and activated by innately attractive stimuli, including odors and taste (fructose). This neuron, therefore, appears to bi-directionally encode stimulus value. Altogether, these findings are in line with our previous studies, where we showed that the MBON-m1 also bi-directionally controls turning behaviour: when it is excited, it inhibits turning and promotes crawling, and when it is inhibited, it promotes turning and inhibits crawling [17].

Our findings suggest that MBON-m1 integrates a variety of sensory information with positive or negative valences, with correspondingly positive or negative weights. The resulting signal loses information about the sensory modality and is classified along a unidimensional valence scale ranging from approach-driving to avoidance-driving cues. Classification along this scale can be made by convergence neurons like MBON-m1 to generate bidirectional signals that are used to control orientation behavior.

## Results

### Cooling sensation pathways from the periphery to the brain

We first investigated the brain pathways downstream of the cooling sensory neurons. As shown previously [11], the three CCs on each side of the head enter the deutocerebrum from the DOG located anteriorly and innervate a region posterior-dorsal relative to the antennal lobe. There, they form an accessory glomerulus juxtaposed to the olfactory glomeruli (**Figure 1A**). The CC synapse onto five cold temperature projection neurons (ctPNs): ctPN1 and 2, further in the text called “Loopers” (very similar to each other based on projection pattern), ctPN3 called “Handlebar”, ctPN4 and ctPN5 (**Figure 1A-C**). Each of these ctPNs receives between 67 and 98 % of their dendritic inputs from the three CCs, with the ctPN1, 2 (Loopers) and 3’s (Handlebar) inputs being almost exclusively from the CCs (respectively 97, 92 and 98%, **Figure 1B**). Considering the similarity of the connection motifs across all CCs (**Figure 1B**), we targeted these cells as a group, using the LexA line for the cooling receptor IR21a (*IR21-LexA)* [24].

**Figure 1.**
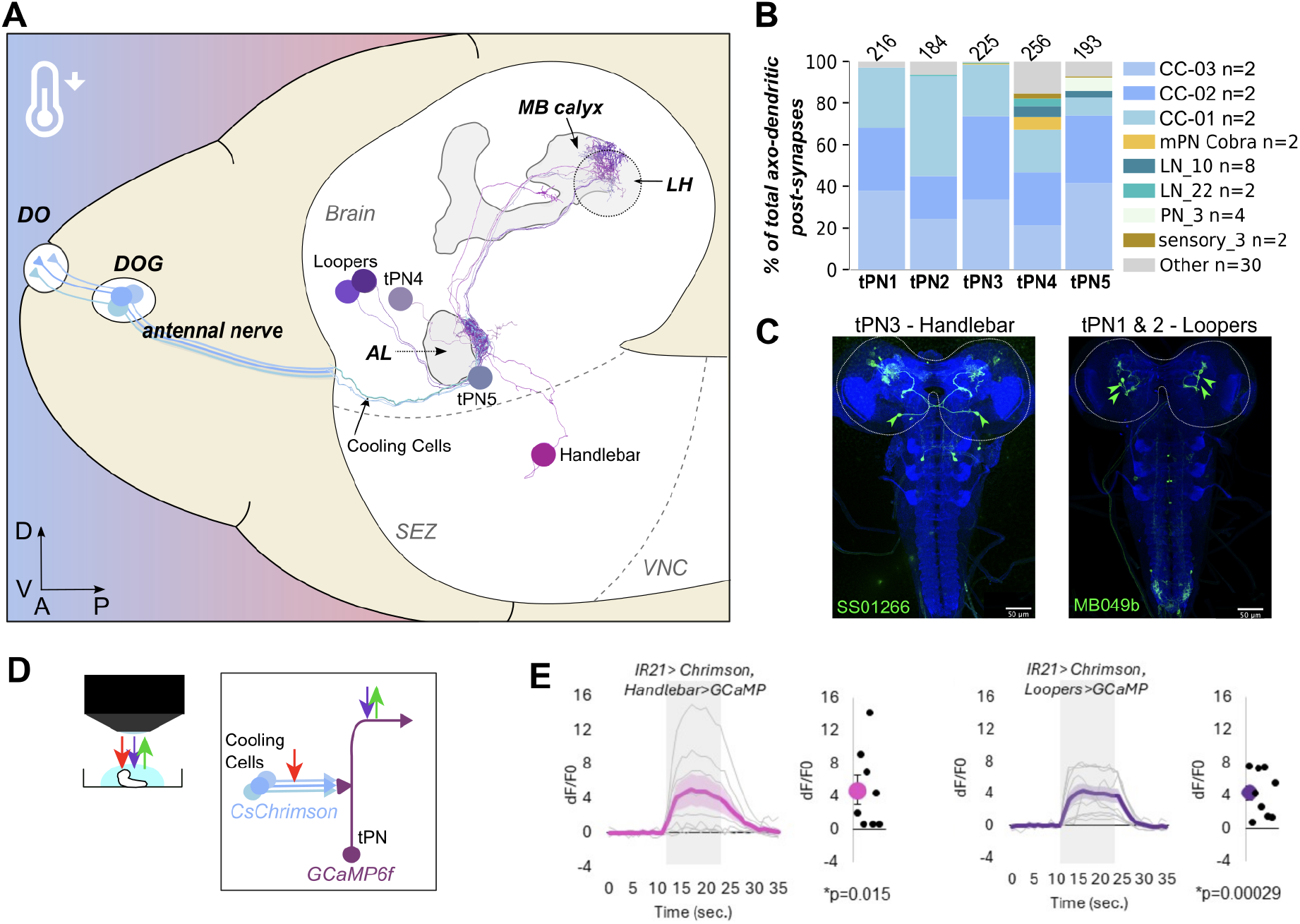
Characterisation of the central circuit for cooling sensation. **a)** EM-based 3D reconstruction of the neural pathway for sensing cooling temperatures: three Cooling-sensing Cells (CCs) located in the dorsal organ (DO) follow the antennal tract and project to a region close to the antennal lobe onto a group of five ctPNs. These neurons send axons towards the higher brain regions spanning from the LH to the MB. **b)** Percent of total synaptic inputs received by the five ctPNs: axo-dendritic inputs are mostly from CCs. The total number of post-synapses considered for each ctPN is shown on top of each bar. See Suppl. Table 1 for details. **c)** We identified the split-GAL4 line SS01266 to target ctPN-3 (Handlebar) and the split-GAL4 line MB049b to target ctPN1-2 (the two Loopers).**d)** To test how the ctPNs respond to CCs activation, we used these lines to drive GCaMP in ctPNs combined with the LexA line IR21a to drive CsChrimson in the CCs. Extracted brains were observed under a 2-photon microscope. We imaged calcium responses in the axonal projections of these neurons. **e)** Calcium transient evoked in the ctPNs in response to CCs optogenetic activation. Both ctPN-3 (Handlebar, N=8 animals) and ctPN1-2 (the Loopers, N=9 animals) showed an excitatory response to CCs’ activation. P-values result from a t-test against chance.

The five identified ctPNs follow slightly different routes, but all target an overlapping, compact brain region that spans the LH and the MB (**Figure 1A**). More specifically, ctPN3 (Handlebar), ctPN4, and ctPN5 closely follow the olfactory uPN tract, while the ctPN1-2 (Loopers’) axons depart from this olfactory bundle and follow the MB peduncle, looping in the ventro-dorsal direction to reach the MB calyces and LH zones (of note, ctPNs with similar anatomical features are also found in the adult *Drosophila*: [25]). ctPN3 (Handlebar) is the only one that projects bilaterally (**Figure 1C**) and strongly contacts the other ctPNs *via* axo-axonic projections (**Suppl. Figure 1**).

We first wanted to verify that ctPNs indeed respond to activation of the CCs. We looked for a genetic line to drive specific expression in some ctPNs and identified a Split-GAL4 line for the ctPN3 (Handlebar, *SS01266*) and another one for ctPN1-2 (both of the two Loopers, *MB049B*) (**Figure 1C**). We expressed CsChrimson in CCs (using *IR21-LexA*) and activated them by a pulse of red light while measuring the intracellular calcium level of ctPN3 (Handlebar) or the ctPN 1-2 (Loopers) using GCaMP6f under a 2-photon microscope (**Figure 1D**). We confirmed that ctPN3 (Handlebar) and ctPN1-2 (Loopers) showed a strong calcium response to the optogenetic activation of the CCs (**Figure 1E**), consistent with the connections observed in the EM volume.

Next, we investigated the behavioral functions of the ctPN3 (Handlebar) and ctPN1-2 (Loopers), especially their role in avoiding cooling temperatures. We silenced one or the other of these ctPN types through constitutive hyperpolarisation *via* Kir2.1 [26]. The larvae were then placed in a linear temperature gradient ranging from an agreeable 21°C to a repulsive 13°C. We found that silencing either type of ctPN significantly impacted larval ability to avoid the low temperature (**Figure 2**).

**Figure 2.**
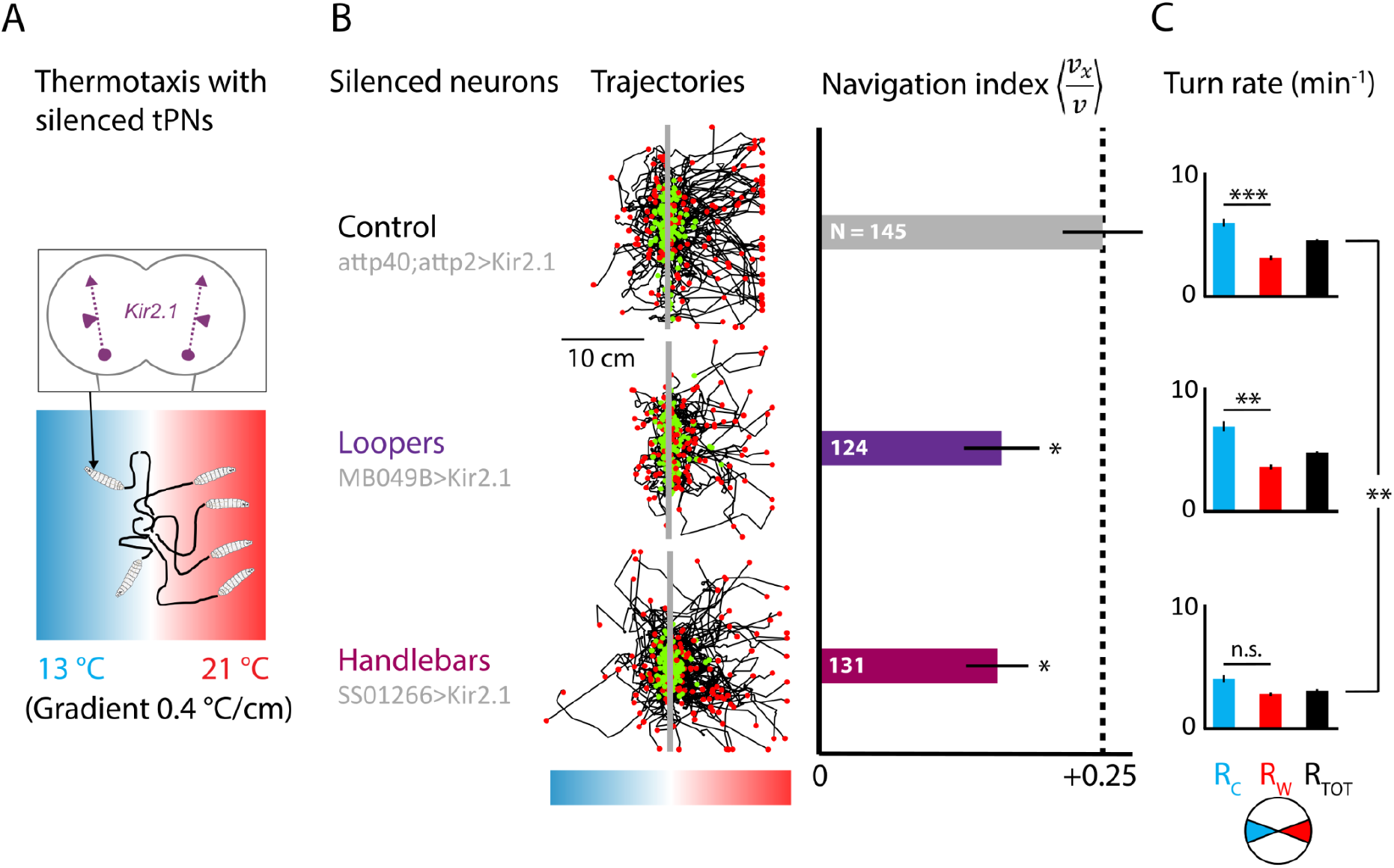
Implication of thermosensory ctPNs in navigation behavior. **a)** CtPNs were silenced with UAS-Kir2.1 and larvae were tested in a 10-minute-long temperature gradient assay. We characterized thermotaxis by **b)** analysing overall trajectories (green circle = start of individual track, red circle = end of track) and computing the navigation index, and **c)** comparing turn rates according to orientation within the gradient (Rc = cooling temperature, Rw = warming temperature). Error bars represent s.e.m. For navigation index, * indicates p<0.05 (Student’s t-test) comparing silenced neuron to control. For turn rate, ** indicates p<0.01, *** indicates p<0.001 (Student’s t-test).

More precisely, the lower navigation index shows that larvae with silenced ctPN3 (Handlebar) or ctPN1-2 (Loopers) distributed more equally in all directions rather than moving more towards the favorable direction, away from cold (**Figure 2B**). The larvae with silenced Handlebar exhibited an overall lower turn rate, and did not show higher turn rate when experiencing a decrease in temperature, compared to an increase, indicating a loss of ability to modulate turn rate in response to changes in temperature (**Figure 2C**). Thus, ctPN3 (Handlebar) and ctPN1-2 (Loopers) relay the aversive signal of cooling temperature detected from the CCs to higher-order brain regions responsible for decision-making. To uncover the circuit logic that integrates this aversive signal with other positive and negative cues, we investigated the downstream targets of ctPNs.

### Thermosensation is relayed to MB and LH where it is mostly segregated from olfaction

As already mentioned, the ctPNs target predominantly two brain areas: the MB and the LH. The different ctPNs send between 0.3 and 25% of their total synaptic projections to three pairs of Kenyon cells (**Figure 3A**) and between 28 and 75% to the LH (**Figure 3B**). The other projections, between 5 and 30%, are mostly onto other PNs, thermosensory, gustatory or multi-glomeruli PNs (**Figure 3C**).

**Figure 3.**
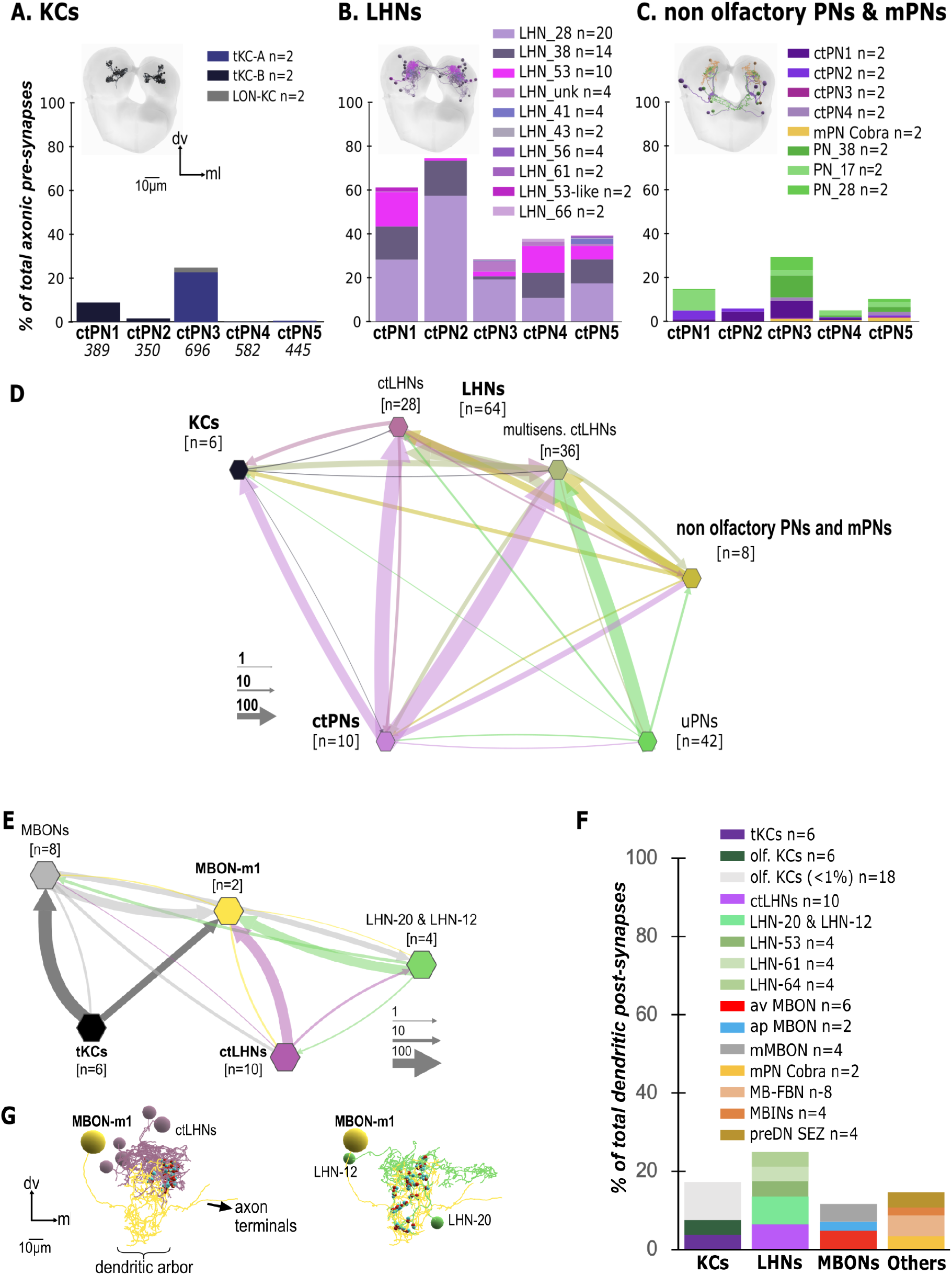
Parallel brain pathways for different sensory modalities converge onto MBON-m1. Bar graphs quantifying the % of synaptic connections between the ctPNs [axonal post-synapses, total n shown under the ctPNs name in **a)**] and the **(a)** MB Kenyon cells, **(b)** LH neurons, and **(c)** projection neurons, including ctPNs, PNs of other modalities, and multiglomeruli PNs (mPNs). LHNs and PNs are clustered according to [27]. Note that the LON-KC receives mostly visual inputs (not shown). The number of neurons in each cluster is shown in brackets. See Suppl. Table 2 with detailed neurons for each cluster. **d)** Connectivity graph showing the connections between the ctPNs and their downstream targets. The olfactory uPNs share common targets with the ctPNs in the LH, these LHNs are grouped under the label “multisensory ctLHNs”. **e)** Connectivity graph showing that MBON-m1 receives, among other, convergent inputs from the tKCs, some of the ctLHPNs, MBONs, and the neurons LHN-20 and LHN-12 (downstream of Or42a and Or42b uPNs, thereby likely with positive valence). **f)** Percent of total synaptic inputs received by MBON-m1 from different neuronal types: KCs, LHNs, MBONs and others. See Suppl. Table 3 with detailed neurons for each cluster. **g)** 3D reconstruction showing the synaptic projections of the ctLHNs (purple) onto MBON-m1 (yellow) and of the olfactory neurons LHN-20 and LHN-12 (green) onto MBON-m1 (yellow). Presynpases are in red and postsynapses in cyan.

Of the two types of KCs targeted by ctPNs, two of them, tKC-A and tKC-B, strictly receive thermosensory inputs (and mPNs inputs for tKC-A), while the third KC, LON-KC, receives mostly visual inputs [21]. One neuron of each of these KCs is found in the right and left MB. Therefore, contrary to the canonical KCs that receive either input from multiple random olfactory PNs or input from a single determined olfactory PN [21], the KCs receiving thermosensory inputs receive inputs from multiple but determined tPNs. More precisely, tKC-A receives massive inputs from ctPN3 (Handlebar), tKC-B receives inputs from ctPN1-2 (both Loopers), and LON-KC receives input only from ctPN3 (Handlebar). To test the effect of signals relayed by the ctPNs onto the MB, we measured the calcium response of the whole population of KCs (using *LexAop-GCaMP6f* under the control of *GMR14H06-LexA*) to the optogenetic activation of ctPN3 (Handlebar) or tPN1-2 (Loopers, using *UAS-CsChrimson* under the control of *SS01266* or *MB049b* [**Suppl Figure 1A]**). The KCs were weakly activated by ctPN3 (Handlebar, [**Suppl Figure 1B]**) and strongly activated by ctPN1-2 (Looper) activation (**Suppl Figure 1C**), suggesting that the ctPNs are excitatory.

In addition, ctPNs target 32 LHNs per brain hemisphere belonging to nine different anatomically defined clusters, according to the classification developed by [27] (**Figure 3B**). Do these LHNs receive mostly thermosensory, rather than olfactory input, like the thermosensory KCs? We addressed this question by asking whether they receive olfactory input via uniglomerular PNs (uPNs] [2,28]). We found that 14 LHNs (per hemisphere) downstream of ctPNs are not targeted by uPNs, whereas 18 integrate uPN and ctPN input (**Figure 3D**). Furthermore, most LHN are also the targets of multi-glomeruli PNs (mPNs) coming from the antennal lobe (**Figure 3D**).

### Downstream of MB and LH, MBON-m1, integrates thermosensory and olfactory information and is required for navigating both sensory gradients

Since the larval thermosensory pathway targets both the LH and the MB, similar to the olfactory pathway [29,30], we looked for downstream partners common to these two sensory systems. In the olfactory system, we described multiple sites of convergence between LHNs and MB neurons involved in ethyl acetate processing, which we previously named ‘convergence neurons’ [17]. We previously found that MBON-m1, one of these neurons, generates a unified, bidirectional instructive output signal by integrating excitatory signals of a positive valence odor from the LH and a mix of excitatory and inhibitory signals from the MB [17]. We found that, in naive animals, the average response of MBON-m1 to an attractive odor was an excitation. However, this response decreased strongly following the formation of an aversive memory for the innately attractive odor [17]. Since MBON-m1 promotes forward crawling when activated and turning when inhibited, these findings align with behavioral observations: naive larvae exhibit odor attraction (correlating with MBON-m1 excitation), while those trained to form an aversive memory show a significant reduction in attraction (correlating with MBON-m1’s decreased response). However, strong avoidance behavior was rarely observed after aversive olfactory learning (at least after 3 training trials used in prior paradigms), nor did we detect any inhibitory response to the aversively conditioned odor.

We therefore wondered whether other sensory cues, in particular more strongly aversive ones that induce active avoidance, could inhibit MBON-m1 and whether this neuron plays a broader role beyond odor processing and more generally participates in the decision to approach or avoid a stimulus according to its valence. To answer these questions, we first used the larval connectome to identify whether MBON-m1 receives synaptic inputs from MB and LHNs downstream of the cooling thermosensory pathways, thought to carry directional signals of negative valence. Indeed, we found that MBON-m1 receives strong inputs from both LH and MB thermosensory pathways (**Figure 3E**).

We also found parallel MB and LH pathways involved in processing innately aversive odor stimuli (e.g. pathways downstream of Or45a/Or47a/Or82a that respond to the repulsive odor geranyl acetate [[2,28]) also reach the MBON-m1, via LHNs, as well as via the KCs. Positive and negative valence odor and aversive thermosensory information are also forwarded directly via mPNs, bypassing the LH and the MB. Thus, MBON-m1 integrates input from multiple appetitive and aversive sensory pathways.

We next tested whether MBON-m1 is also required for navigating positive or negative sensory cues (**Figure 4A-G**). We expressed the inward rectifying channel Kir2.1 in MBON-m1 to render it inactive and tested larval ability to navigate different spatial sensory gradients. We tested larval navigation in cold temperature gradients (decreasing from 21°C), which the larvae are expected to avoid, and in gradient of an attractive odorant ethyl acetate. Silencing MBON-m1 strongly reduced the avoidance of cool temperatures (**Figure 4B**), and we could reproduce the previously shown impairment of ethyl acetate navigation (**Figure 4F**) [17]. Similar to what was observed with larvae in an ethyl acetate gradient, silencing MBON-m1 also increased the overall turn rate (**Figures 4C** and **4G**). MBON-m1 activity thus appears to play a pivotal role in flexible sensory processing and executing behavioral decisions.

**Figure 4.**
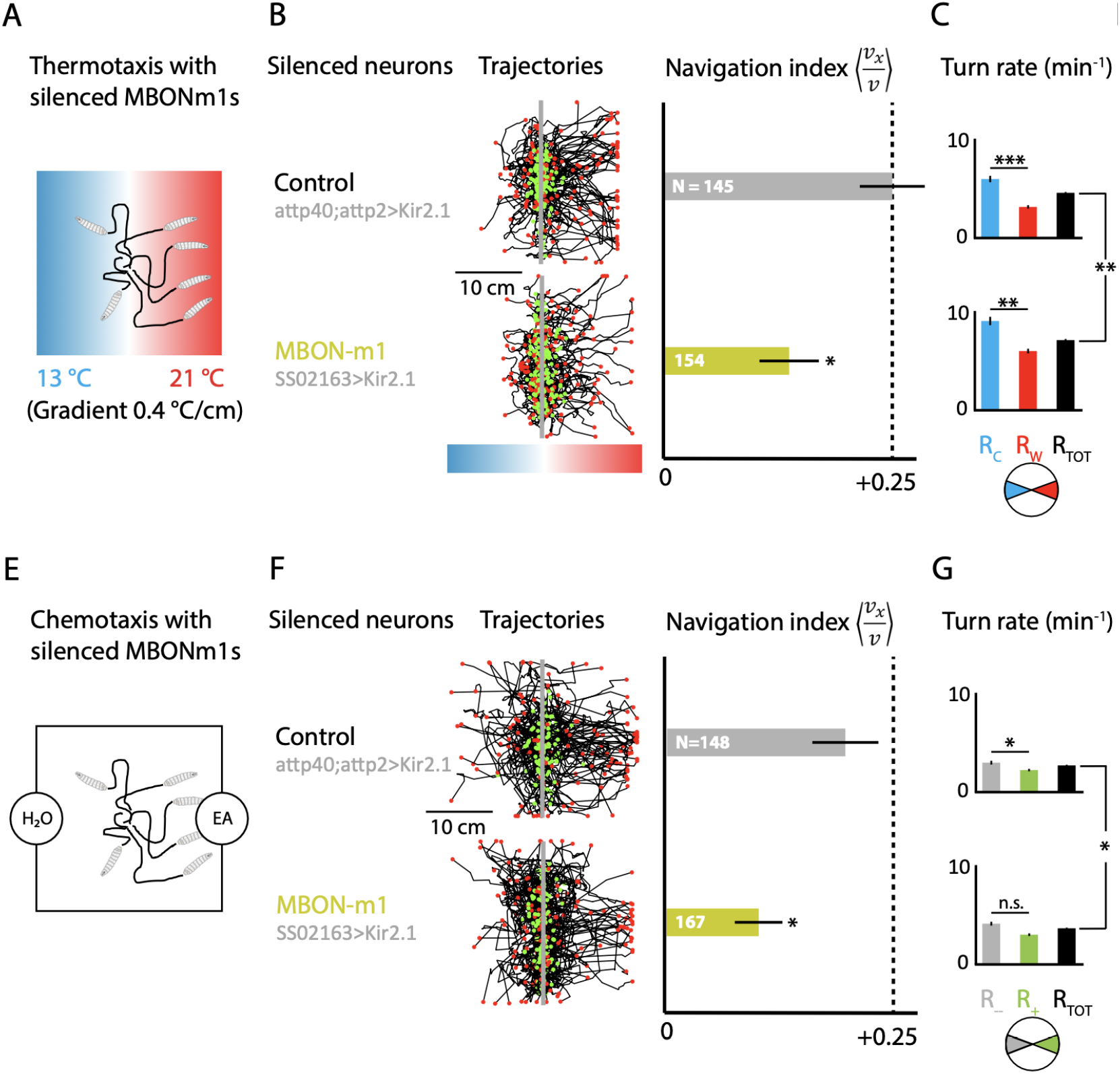
MBON-m1 silencing in thermotaxis and chemotaxis. **a)** SS02163-GAL4 was combined with UAS-Kir2.1 to silence the MBON-m1s in a temperature gradient assay. Larvae were exposed for 10 minutes and we then characterized thermotaxis by computing **b)** the navigation index and recapitulating overall trajectories, **c)** the turn rates according to orientation within the gradient, **d)** the crawl speed of larvae. **e)** The same genetic strategy was used to test the effect of silencing the MBON-m1s in an olfactory gradient assay. We characterized chemotaxis by computing **f)** the navigation index and recapitulating overall trajectories, **g)** the turn rates according to orientation within the gradient, **h)** the crawl speed of larvae. Error bars represent s.e.m. For navigation index, * indicates p<0.05 (Student’s t-test) comparing silenced neuron to control. For turn rate, * indicates p<0.05, ** indicates p<0.01, *** indicates p<0.001 (Student’s t-test) comparing distributions of durations of runs that precede turns.

### MBON-m1 acts as a bidirectional valence integrator - activated by attractants, inhibited by repellents

How are different sensory inputs integrated within MBON-m1, and how do stimuli that drive repulsion affect the activity level of MBON-m1? To answer these questions, we investigated whether we could extend the logic of valence encoding previously described for appetitive odors [17] to other kinds of sensory stimuli (**Figure 5A-C**). We measured the calcium response of MBON-m1 to several distinct innately attractive (ethyl acetate and 1M fructose [**Figure 5B**]), and aversive stimuli (activation of CCs, exposure to geranyl acetate, or 1M salt [**Figure 5C]**). The response of MBON-m1 to these stimuli was bidirectional and followed the valence of the stimuli: The MBON-m1 calcium level decreased in response to sensory cues that usually trigger avoidance (CCs activation, exposure to geranyl acetate or salt [**Fig. 5C**]), whereas it was increased during exposure to attractive stimuli (ethyl acetate or fructose [**Fig. 5B**]). The circuit relaying gustatory cues to the MBON-m1 is currently unknown, because gustatory sensory neurons that project to the brain have not been uniquely identified in this EM volume. In the brain, gustatory input to MBON-m1 could be relayed via LHNs, MBONs, mPNs that receive non-olfactory, potentially gustatory, sensory input [31,32].

**Figure 5.**
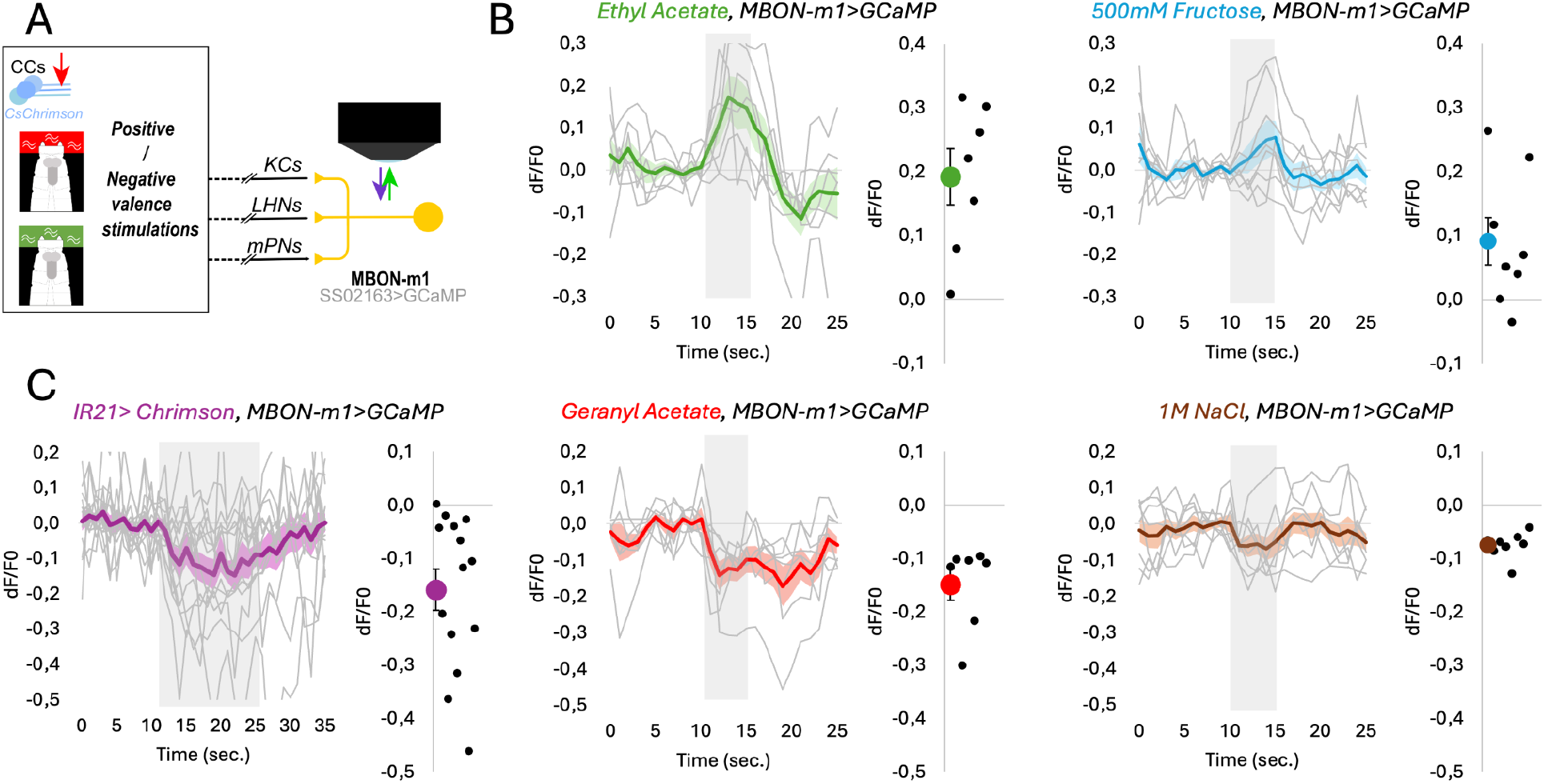
MBON-m1 integrates inputs from diverse sensory modalities and bidirectionally encodes valence. **a)** Stimulation of peripheral sensors either via optogenetic activation (CCs) or application of different odors and tastants via a microfluidics setup with parallel measurement of activity in the MBON-m1 expressing GCaMP. **b)** MBON-m1 is activated by appetitive cues, such as ethyl acetate (N=7 animals) and 500mM Fructose (N=8 animals). **c)** MBON-m1 is inhibited by aversive cues, such as cold cell activation (N=14 animals), geranyl acetate (N=7 animals), and 1M salt (N=8 animals).

The functional imaging results are in line with our behavioral results and the known behavioral effect induced by optogenetic activation of MBON-m1, which promotes approach (represses turn probability) when activated and promotes avoidance (increases turn probability) when inhibited [17]. Therefore, in the presence of attractive cues, MBON-m1 would be activated, which promotes forward crawling towards the cue, while in the presence of aversive stimuli, it would be inhibited, which promotes orientation changes. Based on our functional imaging results, MBON-m1 seems to integrate valence information from multiple sensory cues to inform behavioral decisions.

## Discussion

How positive and negative valence signals of different sensory modalities are integrated and how conflict between them is resolved during action-selection is poorly understood. Here we address this question using the tractable *Drosophila* larva as a model system. In principle, parallel, modality-specific pathways could converge late in the sensory processing hierarchy, at the descending or pre-descending neurons. Alternatively, a coherent valence signal could be computed in higher-order brain areas based on valence information from multiple modalities. To address these questions, we performed an anatomical and functional characterization of the innately aversive cold thermosensory processing pathways and analyzed their patterns of convergence with the previously characterized olfactory pathways.

We first identified thermosensory projection neurons (ctPNs) downstream of the three cool-sensitive sensory cells (CCs) (**Figure 1**) and they are required for normal cool avoidance (**Figure 2**). CtPNs relay temperature information to higher brain centers, including the mushroom body (MB) and the lateral horn (LH), implicated in storing learnt and innate valences of stimuli, respectively (**Figure 3**). We found that these two parallel thermosensory pathways converge again downstream of MB and LH, onto the, so-called, convergence neurons, as previously described for olfactory pathways [17]. Interestingly, we found that the same convergence neuron, MBON-m1, previously implicated in integrating innate and learnt olfactory valences, also integrates innate and learnt thermosensory valences (**Figure 3**). We show that MBON-m1 is involved in navigation in both aversive and appetitive gradients: in avoiding a cool temperature and in approaching an attractive odor (**Figure 4**). This underlines the importance of this single multisensory MBON cell type for larval orientation within various sensory environments. Finally, by imaging calcium signals in MBON-1 in response to distinct appetitive and aversive cues, we that this neuron functions as a multimodal bidirectional valence encoder: it is excited by attractive odors and tastes but inhibited by signals from the cooling pathway and aversive odors or tastes (**Figure 5**). Altogether, our study reveals a multimodal valence integrator that bidirectionally encodes valence and bidirectionally directs behavioral output based on the valence of a sensory cue (**Figure 6**).

**Final 6.**
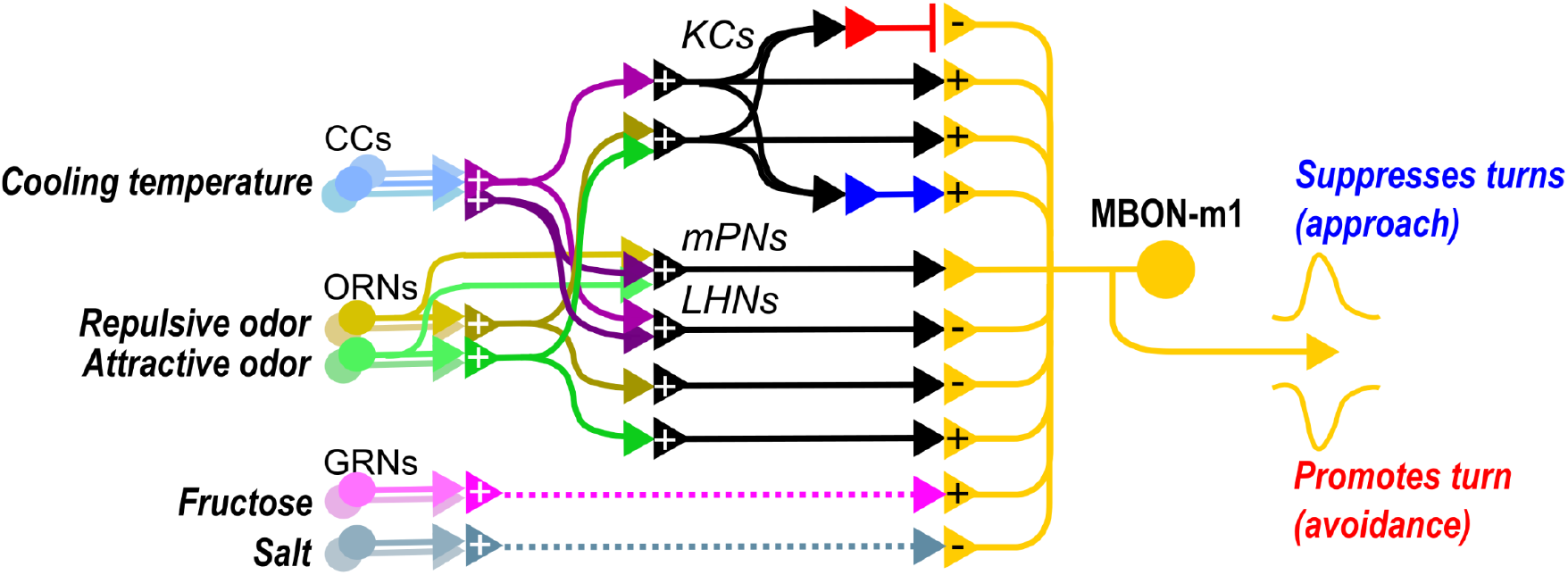
**Schematic representation** of parallel brain pathways for the processing of cooling sensation, attractive odor and aversive odor, showing convergence of these pathways onto MBON-m1. Red and blue arrows depict Mushroom Body output neurons, which relay learned approach and avoidance to the MBON-m1.

### Similarities and differences between thermosensory and olfactory processing circuits

While the sensory neurons for temperature and olfaction exit the dorsal organ via the same tract to reach the anterior part of the brain [11], their projection sites remain spatially segregated. At this point, the excitatory inputs from the three different CCs are simultaneously merged and distributed within five different ctPNs. This differs from the one-to-one connection between OSNs and uPNs in the olfactory system [33] and, similar to the visual system, may enhance the signal-to-noise ratio by pooling the information gathered by the sensors in parallel [16]. Also, unlike the olfactory system, where all uPNs target both Kenyon cells and LH neurons, only some ctPNs (ctPN3 Handlebar and ctPNs1-2 Loopers) strongly project to the MB in addition to the LH, whereas others only project onto LH neurons. If we consider the MB pathway as the main source of behavioral flexibility, this suggests that an important part of the response to cool temperatures is processed independently and potentially differently than in the MB. As ectothermic animals, larvae may rely on a strong innate behavioral component to remain within an optimal temperature range for proper metabolic function [34].

Larvae can learn to approach previously unattractive temperatures if paired with fructose or salt (shown for warmer temperatures, [35]). We found that activating the Handlebar ctPN3 or the Loopers ctPN1-2 induces an excitatory response in the Kenyon Cells, suggesting that MB might also be the site for the formation of such associations, as it is for olfactory associative memories. To go even further on the role of MB in processing odor and temperature cues, an intriguing question is whether KCs representing different sensory modalities interact similarly or differently than the majority of olfactory KCs. In adult flies, KCs typically respond to sensory cues of one modality; however, multisensory associative learning (visual and olfactory) leads to broadening of their response spectra across modalities [36]. There are two KCs per hemisphere, which only receive thermosensory input (tKC-A and tKC-B), however, they form multiple KC-to-KC connections with each other and other KCs with different sensory inputs [37].

### Multiple levels of convergence between olfactory and thermosensory pathways

Our study reveals that olfactory and thermosensory pathways converge at multiple levels within the sensory processing hierarchy. The first site of convergence are multiglomerular PNs (mPNs), which receive direct input from multiple OSNs, as well as from ctPNs. mPNs synapse onto multisensory LHNs, but also form a skip connection that bypasses both the LH and MB, directly to MBON-m1, which might allow the comparison or integration of processed and raw information deeper in the brain [38], and/or may allow specific independent processing of internal state or context information [2]. In the LH, we found a subset of LHNs that selectively receive thermosensory information from ctPNs, but not olfactory information (from uPNs or mPNs). Another subset of multisensory LHNs received input from both ctPNs and uPNs, as well as from mPNs. Finally, these two sensory pathways, converge on convergence neurons, downstream of LH and MB, that integrate learnt and innate valences from both modalities. A multilevel multimodal convergence architecture has been observed for other modalities and has been proposed to be a general feature in multisensory integration circuits, enabling complex response profiles tunable to specific ecological needs [39,40].

### A single neuron bidirectionally encodes valence, independently of sensory modality

We show that MBON-m1 receives convergent inputs from multiple sensory systems via parallel sensory pathways (via LHN, MBINs, KCs, MBONs, mPNs), which carry signals of a specific sensory modality and their innate or learnt valence. Our data suggest that the MBON-m1 pools these sensory signals and extracts the positive *vs*. negative valence information through respectively excitatory *vs*. inhibitory responses to a sensory input. This neuron therefore bidirectionally encodes valence, independently of sensory modality: it is activated by different kinds of attractive cues, and inhibited by different kinds of aversive cues. At the output side, MON-m1 promotes approach if excited and avoidance if inhibited [17]. Our functional imaging data (**Figure 5**) suggest that MBON-m1 is kept at a baseline activity, which can flexibly be pushed towards an increase or decrease in response to different stimuli. Continuous silencing of MBON-m1 using Kir2.1 increases overall turn rate in the presence of both attractive or repulsive stimuli (**Figure 4C** and **G**), which suggests that the baseline activity of MBON-m1 is used to adjust turn rate. Understanding how the activity of this neuron can be maintained in at least three different stable states (baseline, active, and inhibited) is a key future direction and may rely on the highly recurrent loops formed within the MB and the central brain [27,41].

Our findings suggest that a simple integration, linear or not, of all of MBON-m1’s convergent synaptic inputs would result in classifying the sensory cues as “to be approached” or “to be avoided”. Other circuit motives would have allowed such a classification. In particular, negative and positive valence cues could have converged onto different neurons that reciprocally inhibit each other, a motif that is found elsewhere in the central nervous system [14,42] and which allows the selection of mutually exclusive actions. By contrast, the motif of convergent inputs that we describe could favor a graded output, *i*.*e*., going from strong to mild avoidance or mild to strong approach. Indeed, adult and larval *Drosophila* can evaluate the relative value of two stimuli of the same valence [43–45]. Whilst we did not specifically investigate this aspect, the properties of the circuit presynaptic to MBON-m1 suggest that this neuron could encode the value of the stimulus beyond a binary positive or negative valence.

### Bidirectional valence signals guide action-selection and regulate learning

Our findings presented here and in our previous study show that MBON-m1, not only bidirectionally encode valance, but also bidirectionally directs behavioral output based on the valence of a sensory cue [17]: its activation (by appetitive cues) promotes approach and its inhibition below baseline levels of activity (by aversive cues) promotes avoidance. How can activation of a neuron promote one and its inhibition another action? Based on the fact that MBON-m1 is GABAergic [21], we propose that its activation disinhibits one action (crawling), and its inhibition another (turning). Using the available connectome of the *Drosophila* larval brain [27] will enable targeted investigation of circuits downstream of MBON-m1, to reveal how bidirectional valence signals are used to inhibit and disinhibit different combinations of descending neurons, to promote different actions.

While this study focuses on a single CN, MBON-m1, several other CNs integrate thermosensory and olfactory inputs for which genetic tools that would enable their functional investigation do not exist. It is therefore likely that the valence code is distributed across a larger population of CNs. It is interesting to speculate that these different CNs could bidirectionally encode valence in a similar way to MBON-m1, but could control different aspects of the motor response. For example, while MBON-m1 seems to mainly regulate turn vs. crawl behavior during sensory orientation [46] other CNs could modulate other additional behaviors, such as forward vs. backward crawling [47], or rolling [48].

MB output signals also feed back onto the modulatory input of the MB, which is thought to impact memory formation [41]. Specifically, MBON-m1 is connected via a single synaptic step to the appetitive dopaminergic neuron DAN-i1 and the aversive dopaminergic neuron DAN-d1. MBON-m1 activation has been shown to induce an excitatory response in DAN-i1, likely through disinhibition [41]. These feedback signals that encode valence and are directly correlated with the larva’s actions could therefore also influence learning through their effects on dopaminergic neurons. A convergence of movement and reinforcement information in dopaminergic neurons has been observed across the animal kingdom, with important cognitive implications, such as a role in motivational drive.

## Acknowledgements

The authors thank Fly core (especially Monti Mercer) at HHMI Janelia Research Campus, and Nan Hu and Oxana Elliott at the Department of Zoology, University of Cambridge for fly crosses; the Janelia Visitor Project for outstanding support over the years; MRC Laboratory of Molecular Biology Core funding (MZ, AC, KW), HHMI Janelia Research Campus (MZ, AC, CE, BA, MB), Wellcome Trust grant 205038/Z/16/Z (AC), Wellcome Trust grant 205050/Z/16/Z (MZ), ERC grant ERC-2018-COG: 819650 (MZ, CE) for funding. KV was supported by the DFG German Research Foundation (EXC 2117-587 422037984). CE was supported by a Research grant from the Fyssen Foundation and a fellowship from the ATIP-Avenir Program.

## Material & Methods

### Fly husbandry and fly lines

#### Functional imaging (Microfluidics)

For larval experiments not involving optogenetics, flies were reared at 22°C under a 12-hour light/12-hour dark cycle and 60% relative humidity in vials containing standard cornmeal agar–based medium. Adult flies were transferred to larvae collection cages (Genesee Scientific) containing grape juice agar plates and 180 mg of fresh yeast paste per cage. Flies were allowed to lay eggs on the agar plate for 1 to 2 days before the plate was removed for the collection of larvae in the different developmental stages. Calcium imaging experiments and anatomical studies were performed in L1 larvae (2 days AEL).

#### Functional imaging (Optogenetics)

Larvae were reared in the dark at 25°C in food vials. The food was supplemented with trans-retinal (SIGMA R2500) at a final concentration of 500μM. For imaging experiments, the larvae were selected at their first-instar stage. To assess the response to optogenetic activation of CCs, IR21a-LexA was crossed to direct the expression of LexAop-Chrimson.

#### Behavior experiments

Larvae were reared at room temperature (approximately 23-24°C) in collection cages (Gensee) with grape juice plates with yeast paste added. Plates were transferred every 24 hours, and early second-instar larvae were selected for experiments, rinsed three times in DI water, and then transferred to a plain agar plate for 5 minutes before experimental testing. The empty stock y w;attP40;attP2 was crossed to the effector as a baseline control.

#### Fly lines

The following fly lines were used:

UAS-GCaMP6m; Orco::RFP (Harvard) - UAS-GCaMP6m, BDSC #42748; Orco::RFP, BDSC #63045); 20XUAS-CsChrimson-mVenus 53 - BDSC #55134; 13XLexAop2-CsChrimson-tdTomato in attP18; Ir21a-LexAp65 in JK22C;20XUAS-IVS-GCaMP6f 15.693 in VK00005; UAS-Kir2.1 - BDSC #6596; Empty GAL4: y w;attP40;attP2 - BDSC #79603; CCs - IR21a-LexA; KCs -14H06-LexA; ctPN1-2: Loopers - MB049b: w[1118]; GMR60H12-p65.AD; GMR14E09-GAL4.DBD - BDSC #603334;ctPN3: Handlebar - SS01266: GMR_43A02_XA_21-x-GMR_20F01_XD_01; MBON-m1 - SS02163: GMR_52H01_XA_21-x-GMR_40F09_XD_01 - BDSC #604151

### Experimental Assays

#### Behavior

For thermotaxis experiments, the arena housed a 1D linear temperature gradient platform. As previously described in [11], cold and warm reservoirs were maintained with thermoelectric coolers (TECs) under PID control, with liquid cooling blocks dissipating excess heat. A 6-mm-thick rectangular aluminum sheet bridged the two reservoirs and established a stable linear gradient. A 3-mm thick, 22 × 22 cm agar gel (2.5% wt./vol. agar, 0.75% wt./vol. charcoal) was placed on the rectangular sheet. Approximately 20 larvae were placed on the gradient for each trial, free to crawl for 10 minutes on the thermal gradient gel (13 C on the cold side, 21 C on the warm side). The arena was illuminated using strips of red LEDs (640 nm, outside the visible spectrum for larvae), and behavior was recorded with an above-mounted CCD camera (Basler). The position and body contour of each larva was determined using the MAGAT Analyzer software [10], then individual larva tracks were segmented into forward-crawling runs and direction-altering reorientations. Other Matlab scripts sorted the crawling data into properties of each run, and software written in Igor Pro calculated relevant behavior metrics, such as navigation index, turning rates, etc. [10]. Same thermotaxis data for empty-GAL4/UAS-kir control is shown in **Figures 2** and **4**.

For chemotaxis experiments, experiments were performed in the same arena and with the same protocol and analysis as for thermotaxis, but with temperature controllers turned off, and with plastic dishes with lids. Ethyl Acetate was obtained from Sigma-Aldrich and used at the indicated dilution (EA, 10−4 (CAS #141-78-6)). Before each experiment, a 12-mm plastic cup at one edge was loaded with 200 μl of odorant solution.

#### Functional imaging (Microfluidics)

We used a previously described method for microfluidic delivery of odorants in aqueous form with simultaneous imaging of calcium activity in intact larvae [2,28,46]. All experiments used an eight-channel microfluidic chip equipped with a vacuum port to stabilize the animal’s head. The same odorants were used as in the behavioral experiments, but at lower concentrations (GA = 10^−6^; EA = 10^−5^). Tastants were also diluted (Fructose = 500 mM (CAS# 57-48-7); NaCl = 1M (CAS# 7647-14-5) [49]. Stimuli consisted of 5-s pulses interleaved with 15-s water washout periods. An L1 larva was washed with DI water and loaded into the microfluidic device using a 1-ml syringe filled with Triton X-100 [0.1% (v/v)] solution. The animal was pushed to the end of the loading channel, with its dorsal side facing the objective. GCaMP signal was recorded using an inverted Nikon Ti-E spinning disc confocal microscope and a 60× water immersion objective [numerical aperture (NA), 1.2]. A CCD microscope camera (Andor iXon EMCCD) captured frames at 30 Hz. Recordings from 7-8 larvae were collected for each condition. The same larvae were tested for all odors (N=7), and a different set of larvae was tested for all tastants (N=8).

#### Functional imaging (Optogenetics)

Central nervous systems were dissected in a cold buffer containing 135 mM NaCl, 5 mM KCl, 4 mM MgCl2·6H2O, 2 mM CaCl2·2H2O, 5 mM TES and 36 mM sucrose, pH 7.15 (Marley and Baines, 2011) and adhered to poly-L-lysine (SIGMA, P1524) coated cover glass in small Sylgard (Dow Corning) plates. Optogenetic activation was done by red flood illumination on the sample (625 nm, Four channel LED driver, Thorlabs, power) through the objective. Light stimulations were delivered for 3 s and for four successive times (ISI *ca*. 15 s) in each scan. Larvae were imaged at 35 fps using a multiphoton microscope equipped with a fast resonant galvo scan module (customized Bergamo Multiphoton, Thorlabs) controlled by ScanImage 2016 (http://www.scanimage.org). The light source was a femtosecond pulsed laser tuned to 925 nm (Mai Tai, Spectraphysics). The objective was a 25 X water immersion objective (NA 1.1 and 2 mm WD, Nikon). For image analysis, image data were processed using custom code in Matlab (The Mathworks, Inc). Specifically, the code automatically corrects for misaligned images, determines the regions of interest (ROIs) from maximum intensity projection of entire time series images, and measures the mean intensity of the ROIs minus the background fluorescence. In all cases, changes in fluorescence were calculated relative to baseline fluorescence levels (*F*0) as determined by averaging over a period of 5 s. just before the optogenetic stimulation. The fluorescence values were calculated as (*F***t** - *F***0**)/*F***0**, where *F***t** is the fluorescent mean value of a ROI in a given frame. Analyses were performed on the average of the consecutive four stimulations.

### Statistics

#### Behavior

Behavioral larva data were filtered to only include tracks that started during the first 200 seconds of the experiment and lasted at least 200 seconds – this prevented any double counting, so every larva was associated with only a single track. The navigation index was calculated as the average x-direction velocity during runs <v_x>, divided by the average speed during runs <v>. The number was computed for each individual, and the overall navigation index (NI) was the average of the individual NI, and error bars were calculated as the standard error of the mean in the list of NI values. Significance comparisons were calculated comparing lists of NI values, using Student’s t-test to determine the p value. Turning rates were determined by dividing runs into octants of crawling direction, and computing R_oct = N_oct/T_oct (R the turning rate, N the number of turns performed, T the total time spent crawling in that direction).

#### Functional Imaging

For each larva, the peak response during stimulus presentation was compared to the chance level, analyzed via a normal t-test. Responses during stimulation were normalized based on the baseline response 5s before the stimulation. P-values for each experiment are given in the respective figure panel. Each graph shows the mean calcium signal plotted as the relative response strength ΔF/F and the related standard error of the mean on the y-axis. The time in seconds is given below each graph on the x-axis. The grey box indicates the duration of the stimulus application. The sample size of each group (N=7-8) is given above each row. n.s. p > 0.05; * p < 0.05. Comparisons of before *vs*. during stimulation were done using a t-test against zero on the normalised data.

