## Supplementary Information for "A multisensory, bidirectional, valence encoder guides behavioral decisions"

#### Figures

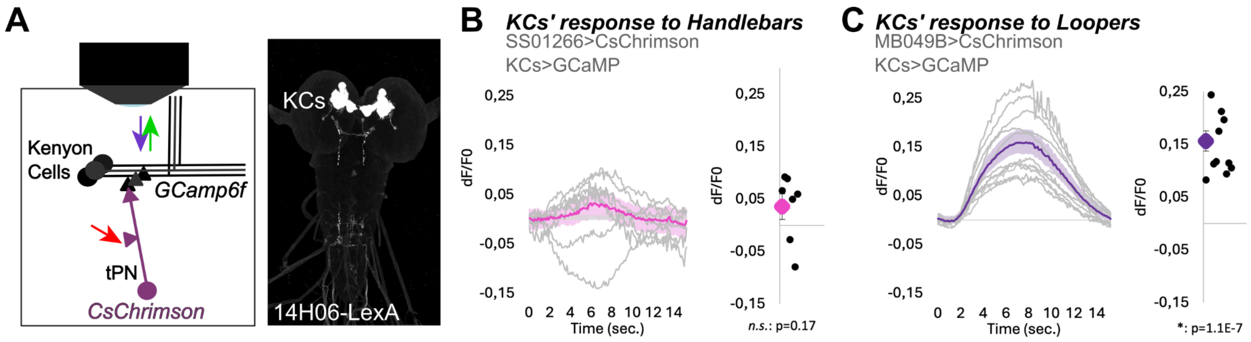


**Supplementary Figure 1 associated with Figure 3.**
**a)** To test the input signs of ctPN1-2 (Loopers) and ctPN3 (Handlebar) onto the KCs, we measured the calcium response of the whole population of KCs (using the 14H06-LexAline and LexAop-GCamp6f) to the optogenetic activation of these ctPNs. **b)** We used the split-GAL4 line SS01266 and UAS-CsChrimson to optogenetically activate ctPN3 Handlebar and recorded the calcium response of KCs, revealing mostly a weak activation of the KCs to ctPN3 Handlebar input, more rarely an inhibitory response. The averaged response of KCs to ctPN3 activation is not significantly different from zero (**N=7 animals**). **c)** We used the same approach to record the response of KCs to the optogenetic activation of the ctPN1-2 Loopers using MB049b split-GAL4 line to activate these thermo-PNs (**N=11 animals**). Here, Kenyon Cells are strongly activated by ctPN1-2 Loopers stimulation.

### Tables

**Supplementary Table 1**

List of the neurons presynaptic to the five ctPNs (dendrite). Only the neurons whose inputs represent >1% of the total synaptic inputs of these neurons are shown. Weaker synaptic inputs are grouped as “Others”.

| ID Left neuron | ID Right neuron | Celltype | MW_level_7_cluster | Group name | tPN4 (suckerfish) | tPN3 (Handle bar) | tPN1 (Lower Looper) | tPN2 (Upper Looper) | tPN5 (Only connects) | TotSyn |
|---|---|---|---|---|---|---|---|---|---|---|
| 3609202 | 3487306 | sensory | 5 | CC-3 | 55 | 76 | 82 | 45 | 80 | 408 |
| 3639968 | 18630153 | sensory | 5 | CC-2 | 65 | 90 | 65 | 38 | 63 | 381 |
| 3608397 | 3650600 | sensory | 5 | CC-1 | 52 | 56 | 63 | 88 | 17 | 326 |
| 16795838 | 16795524 | PN | 10 | mPN Cobra | 16 | 1 | 0 | 0 | 0 | 689 |
| 8102935 | 7600311 | LN | 10 | LN_10 | 7 | 0 | 0 | 0 | 1 | 306 |
| 8274021 | 11051276 | LN | 10 | LN_10 | 0 | 0 | 0 | 0 | 5 | 459 |
| 8877971 | 3806573 | LN | 10 | LN_10 | 3 | 0 | 0 | 0 | 0 | 377 |
| 7939890 | 6557581 | LN | 10 | LN_10 | 3 | 0 | 0 | 0 | 0 | 1107 |
| 17844900 | 18598859 | LN | 22 | LN_22 | 10 | 1 | 0 | 1 | 0 | 231 |
| 11821320 | 6220384 | PN | 3 | PN_3 | 0 | 0 | 0 | 0 | 8 | 174 |
| 5327961 | 11184236 | PN | 3 | PN_3 | 0 | 0 | 0 | 0 | 4 | 217 |
| 4338596 | 15652529 | sensory | 3 | sensory_3 | 6 | 0 | 0 | 0 | 1 | 364 |
| Other | | | | Other | 39 | 1 | 6 | 12 | 14 | |
| **Total synapses** | | | | | **256** | **225** | **216** | **184** | **193** | |

**Supplementary Table 2**

List of the neurons postsynaptic to ctPNs (axonal part). Only the neurons receiving >1% of their total synaptic inputs from the ctPNs are shown.

| ID Left neuron | ID Right neuron | Cell type | MW_level_7_cluster | Group name | tPN4 | tPN3 Handlebar | tPN1 Lower Looper | tPN2 Upper Looper | tPN5 | Total Syn | sum | total input >1% |
|---|---|---|---|---|---|---|---|---|---|---|---|---|
| 4279887 | 10293348 | LHN | 28 | **LHN_28** | 8 | 44 | 0 | 0 | 9 | 207 | 61 | Y |
| 9747418 | 4127707 | LHN | 28 | **LHN_28** | 28 | 59 | 1 | 3 | 10 | 196 | 101 | Y |
| 4635057 | 7971031 | LHN | 28 | **LHN_28** | 0 | 3 | 23 | 22 | 7 | 148 | 55 | Y |
| 19611917 | 8138616 | LHN | 28 | **LHN_28** | 1 | 1 | 15 | 28 | 1 | 200 | 46 | Y |
| 12745771 | 2914206 | LHN | 28 | **LHN_28** | 2 | 3 | 18 | 44 | 2 | 233 | 69 | Y |
| 7582412 | 2776773 | LN/LHN | 28 | **LHN_28** | 16 | 2 | 11 | 2 | 11 | 141 | 42 | Y |
| 4570270 | 2921933 | LN/LHN | 28 | **LHN_28** | 3 | 8 | 7 | 14 | 11 | 189 | 43 | Y |
| 12948470 | 9275210 | LN/LHN | 28 | **LHN_28** | 4 | 4 | 30 | 50 | 3 | 158 | 91 | Y |
| 4636432 | 9808146 | LN/LHN | 28 | **LHN_28** | 1 | 10 | 5 | 38 | 17 | 148 | 71 | Y |
| 12387154 | 8723267 | LHN | 38 | **LHN_38** | 16 | 0 | 0 | 0 | 2 | 192 | 18 | Y |
| 9201911 | 17591687 | LHN | 38 | **LHN_38** | 9 | 0 | 0 | 0 | 14 | 70 | 23 | Y |
| 16474159 | 9291437 | LHN | 38 | **LHN_38** | 6 | 10 | 0 | 0 | 8 | 79 | 24 | Y |
| 9421292 | 5507096 | LHN | 38 | **LHN_38** | 4 | 0 | 53 | 7 | 2 | 173 | 66 | Y |
| 17420035 | 8705088 | LHN | 38 | **LHN_38** | 14 | 0 | 0 | 0 | 1 | 81 | 15 | Y |
| 4778745 | 4319395 | LHN | 38 | **LHN_38** | 16 | 0 | 0 | 0 | 18 | 94 | 34 | Y |
| 9445876 | 4102413 | LN/LHN | 38 | **LHN_38** | 2 | 0 | 6 | 49 | 4 | 367 | 61 | Y |
| 4379603 | 8307514 | LHN | 53 | **LHN_53** | 0 | 0 | 0 | 0 | 5 | 206 | 5 | Y |
| 4391317 | 11826171 | LHN | 53 | **LHN_53** | 18 | 13 | 0 | 0 | 21 | 213 | 52 | Y |
| 9207332 | 8305958 | LHN | 53 | **LHN_53** | 11 | 0 | 0 | 0 | 0 | 156 | 11 | Y |
| 7582489 | 7977834 | LHN | 53 | **LHN_53** | 1 | 2 | 61 | 4 | 0 | 185 | 68 | Y |
| 2439134 | 4099900 | LHN | 53 | **LHN_53** | 41 | 0 | 0 | 0 | 1 | 316 | 42 | Y |
| 10540188 | 8932110 | LHN | NA | **LHN_unk** | 6 | 29 | 1 | 0 | 1 | 218 | 37 | Y |
| 11123016 | 4152020 | LHN | NA | **LHN_unk** | 6 | 4 | 0 | 0 | 3 | 67 | 13 | Y |
| 4780707 | 15683275 | LHN | 41 | **LHN_41** | 28 | 2 | 0 | 0 | 5 | 294 | 35 | Y |

| | | | | | | | | | | | | |
|---|---|---|---|---|---|---|---|---|---|---|---|---|
| 11135340 | 11645541 | LN/LHN | 41 | **LHN_41** | 0 | 0 | 0 | 0 | 6 | 283 | 6 | Y |
| 5118060 | 15605987 | LHN | 43 | **LHN_43** | 30 | 3 | 0 | 0 | 1 | 760 | 34 | Y |
| 4379508 | 3794023 | LHN | 56 | **LHN_56** | 12 | 0 | 0 | 0 | 2 | 132 | 14 | Y |
| 4785242 | 2935198 | LN/LHN | 56 | **LHN_56** | 6 | 0 | 0 | 0 | 1 | 182 | 7 | Y |
| 16442113 | 3814175 | LHN | 61 | **LHN_61** | 10 | 0 | 0 | 0 | 1 | 152 | 11 | Y |
| 9287199 | 4607160 | LHN | NA | **LHN_53-like** | 1 | 1 | 7 | 1 | 1 | 116 | 11 | Y |
| 8118474 | 4229950 | KC | 40 | **tKC-A** | 1 | 159 | 0 | 0 | 3 | 692 | 163 | Y |
| 8262372 | 8068173 | KC | 40 | **LON-KC** | 1 | 15 | 0 | 0 | 0 | 567 | 16 | Y |
| 7187384 | 3664102 | KC | 40 | **tKC-B** | 0 | 0 | 35 | 6 | 0 | 469 | 41 | Y |
| 16795838 | 16795524 | PN | 10 | **mPN Cobra** | 4 | 9 | 0 | 0 | 8 | 542 | 21 | Y |
| 12156009 | 7020344 | PN | 17 | **ctPN3** | 0 | 2 | 0 | 0 | 2 | 468 | 4 | Y |
| 8293958 | 3869488 | PN | 17 | **ctPN1** | 6 | 54 | 4 | 16 | 0 | 304 | 80 | Y |
| 11146422 | 19276024 | PN | 25 | **ctPN2** | 2 | 0 | 16 | 5 | 3 | 223 | 26 | Y |
| 4108016 | 3002410 | PN | 28 | **ctPN4** | 0 | 12 | 0 | 0 | 7 | 337 | 19 | Y |
| 3991518 | 9747710 | PN | 38 | **PN_38** | 5 | 70 | 0 | 0 | 10 | 269 | 85 | Y |
| no pair | 4620453 | PN | 17 | **PN_17** | 1 | 4 | 7 | 0 | 4 | 37 | 16 | Y |
| 4985759 | 9291474 | PN | 17 | **PN_17** | 12 | 12 | 29 | 0 | 6 | 109 | 59 | Y |
| 4154421 | 3690921 | PN | 28 | **PN_28** | 0 | 43 | 2 | 0 | 6 | 110 | 51 | Y |
| 5091874 | 9287906 | LHN | 66 | **LHN_66** | 7 | 0 | 0 | 0 | 0 | 442 | 7 | Y |
| 5959820 | 8943485 | LHN | 28 | **LHN_28** | 0 | 0 | 0 | 0 | 7 | 64 | 7 | Y |
| Other | | | | **Other** | 243 | 118 | 58 | 61 | 221 | | | N |
| Total synapses | | | | | 582 | 696 | 389 | 350 | 445 | | | |

**Supplementary Table 3**

List of the neurons presynaptic to MBON-m1 (dendrite). Only the neurons whose inputs represent >1% of the total synaptic inputs of MBON-m1 are shown.

| ID Left neuron | ID Right neuron | Cell type | Group name | MW_level_7_cluster | MBON_m1 | DS of ctPNs | total input >1% |
|---|---|---|---|---|---|---|---|
| | | | | **total synapses:** | **1613** | | |
| 8509534 | 16515871 | PN | **mPN cobra** | 29 | 54 | N | **Y** |
| 8262302 | 5051342 | KC | **olf KC** | 30 | 20 | | **Y** |
| 5937084 | 8850802 | KC | **olf KC** | 30 | 21 | | **Y** |
| 4117305 | 16720240 | KC | **olf KC** | 36 | 20 | | **Y** |
| 8118474 | 4229950 | KC | **tKC** | 40 | 28 | Y | **Y** |
| 8262372 | 8068173 | KC | **tKC** | 40 | 20 | Y | **Y** |
| 4379603 | 8307514 | LHN | **ctLHN** | 53 | 30 | Y | **Y** |
| 9207332 | 8305958 | LHN | **ctLHN** | 53 | 21 | Y | **Y** |
| 7582489 | 7977834 | LHN | **ctLHN** | 53 | 29 | Y | **Y** |
| 16386470 | 17013169 | LHN | **LHN_53** | 53 | 47 | | **Y** |
| 8798010 | 4195012 | MBON | **av MBON** | 59 | 20 | | **Y** |
| 8338584 | 4230749 | MBON | **av MBON** | 59 | 25 | | **Y** |
| 7802210 | 16797672 | MBON | **av MBON** | 59 | 31 | | **Y** |
| 5938794 | 8310955 | LHN | **LHN_61** | 61 | 36 | | **Y** |
| 15766806 | 13863331 | LHN/MB-FBN | **LHN-20 & LHN-20** | 61 | 31 | | **Y** |
| 16089743 | 10340880 | LHN | **LHN_61** | 61 | 23 | | **Y** |
| 14082322 | 7840791 | MBON | **ap MBON** | 64 | 40 | | **Y** |
| 15398730 | 6703240 | MBON | **mMBON** | 64 | 33 | | **Y** |
| 12832017 | 19091931 | MB-FFN | **LHN_64** | 64 | 35 | | **Y** |
| 11257084 | 7612147 | LHN | **LHN-20 & LHN-20** | 64 | 83 | | **Y** |
| 10543751 | 11464638 | CN | **LHN_64** | 64 | 26 | | **Y** |
| 15594113 | 15590555 | pre-DN-SEZ | **pre-DN-SEZ** | 76 | 61 | | **Y** |
| 4408397 | 4199033 | MB-FBN | **MB-FBN** | 81 | 49 | | **Y** |
| 9062992 | 8792477 | MB-FBN | **MB-FBN** | 81 | 18 | | **Y** |

| | | | | | | | |
|---|---|---|---|---|---|---|---|
| 9064208 | 5934511 | MB-FBN | **MB-FBN** | 81 | 19 | | **Y** |
| 11017761 | 16178283 | MBON | **mMBON** | 82 | 40 | | **Y** |
| 9287199 | 4607160 | n | **ctLHN** | NA | 22 | Y | **Y** |
| 4391317 | 11826171 | LHN | **ctLHN** | 53 | 17 | Y | **Y** |
| 12138418 | 8310087 | LHN | **LHN_53** | 53 | 17 | | **Y** |
| 4381377 | 10673895 | MBIN | **MBIN** | 57 | 17 | | **Y** |
| 4381129 | 15404207 | MBIN | **MBIN** | 57 | 17 | | **Y** |
| 18500988 | 14334367 | pre-DN-SEZ | **pre-DN-SEZ** | 81 | 18 | | **Y** |
| 17177005 | 8019816 | MB-FBN | **MB-FBN** | 81 | 27 | | **Y** |
